# Balancing positive and negative selection: *in vivo* evolution of *Candida lusitaniae MRR1*

**DOI:** 10.1101/2020.11.30.405712

**Authors:** Elora G. Demers, Jason Stajich, Alix Ashare, Patricia Occhipinti, Deborah A. Hogan

## Abstract

The evolution of pathogens in response to selective pressures present during chronic infections can influence persistence, virulence, and the outcomes of antimicrobial therapy. Because subpopulations within an infection can be spatially separated and the host environment can fluctuate, an appreciation of the pathways under selection may be most easily revealed through the analysis of numerous isolates from single infections. Here, we continued our analysis of a set of clonally-derived *Clavispora (Candida) lusitaniae* isolates from a single chronic lung infection with a striking enrichment in the number of alleles of *MRR1*. Genetic and genomic analyses found evidence for repeated acquisition of gain-of-function mutations that conferred constitutive Mrr1 activity. In the same population, there were multiple alleles with both gain-of-function mutations and secondary suppressor mutations that either attenuated or abolished the constitutive activity suggesting the presence of counteracting selective pressures. Our studies demonstrated tradeoffs between high Mrr1 activity, which confers resistance to the antifungal fluconazole, host factors, and bacterial products through its regulation of *MDR1*, and resistance to hydrogen peroxide, a reactive oxygen species produced in the neutrophilic environment associated with this infection. This inverse correlation between high Mrr1 activity and hydrogen peroxide resistance was observed in multiple *Candida* species and in serial analysis of populations from this individual collected over three years. These data lead us to propose that dynamic or variable selective pressures can be reflected in population genomics and that these dynamics can complicate the drug resistance profile of the population.

**Importance:** Understanding microbial evolution within patients is critical for managing chronic infections and understanding host-pathogen interactions. Here, our analysis of multiple *MRR1* alleles in isolates from a single *Clavispora (Candida) lusitaniae* infection revealed the selection for both high and low Mrr1 activity. Our studies reveal tradeoffs between high Mrr1 activity, which confers resistance to the commonly used antifungal fluconazole, host antimicrobial peptides and bacterial products, and resistance to hydrogen peroxide. This work suggests that spatial or temporal differences within chronic infections can support a large amount of dynamic and parallel evolution, and that Mrr1 activity is under both positive and negative selective pressure to balance different traits that are important for microbial survival.

## Introduction

Understanding the positive and negative selective pressures that shape drug resistance profiles in microbial populations is critical for combating the development of antimicrobial resistance, an ever-increasing problem in clinical settings. Increased drug resistance in bacteria and fungi has been associated with clinically- and agriculturally-used antimicrobial agents (reviewed in (1–3)), and drug resistance elements may be selected for based on their ability to protect against factors produced by other microbes or plant, animal, and insect hosts (4, 5). Based on the analysis of bacterial isolates of *Burkholderia dolosa* or *Pseudomonas aeruginosa* from single patients and across cohorts of patients, it is clear that *in vivo* factors can lead to the repeated selection for subpopulations with the same genes or pathways mutated (6–8). Furthermore, there is evidence that pathways can be upregulated then downregulated in the same phylogenetic lineages. For example, suppressor mutations within *P. aeruginosa algU* frequently arise in strains with high AlgU signaling caused by mutations in the gene encoding the AlgU repressor MucA (9). Less is known about the negative selective pressures acting against sustained microbial resistance.

In Demers *et al.* (10), we described a set of twenty recently-diverged *Clavispora* (*Candida*) *lusitaniae* isolates obtained from the lung infection of a single individual with cystic fibrosis (CF). *C. lusitaniae* is among the emerging non-*albicans Candida* spp. that cause life threatening disseminated infections in individuals who are immunocompromised (11, 12), and can cause infections of the gastrointestinal tract (13–15), surgical sites, or implanted devices in immunocompetent individuals. *C. lusitaniae* is notorious for its rapid development of resistance to antifungal drugs including amphotericin B, azoles and echinocandins (14, 16–19) and, relative to *Candida albicans* and other *Candida* species that are both opportunistic pathogens and members of the mycobiome, it is more closely related to *Candida auris*, a species in which multi-drug resistant isolates have caused hospital-associated outbreaks (20–26). Our previous analyses of heterogeneity in fluconazole (FLZ) resistance among these isolates identified numerous distinct alleles of *MRR1* (*CLUG_00542*). Multiple alleles encoded gain-of-function (GOF) mutations causing constitutive Mrr1 activity, which, as in other *Candida* species, increased expression of *MDR1* and Mdr1 multidrug efflux pump activity (10, 27–32). At the time that these isolates were recovered, the patient had no history of antifungal treatment, suggesting that selection for constitutively active Mrr1 variants may have been driven by the need for resistance to other host- or microbe-produced compounds. Within this study, however, we found multiple lineages with recently evolved *MRR1* alleles that rendered cells more sensitive to FLZ than even *mrr1*Δ strains. Here, we address the perplexing question of why this population had recently diverged *MRR1* alleles that encoded both high and low Mrr1 activity. To do so, we expressed both native and synthesized *MRR1* alleles that represent intermediates during *MRR1* evolution in a common genetic background and tested the effects of these alleles on growth in *in vivo* relevant conditions. Using genetics and genomics, we concluded that multiple *C. lusitaniae MRR1* alleles that conferred low Mrr1 activity resulted from initial mutation that caused constitutive Mrr1 activity followed by a second mutation that either suppressed constitutive activation or inactivated the protein. Constitutive Mrr1 activity caused increased sensitivity to a variety of biologically relevant compounds including hydrogen peroxide (H_2_O_2_) and suppression of constitutive Mrr1 activity rescued growth under many of these conditions. Monitoring populations from respiratory samples from this subject over time supports the model that there are opposing selective pressures *in vivo* that select for and against constitutive Mrr1 activity, as reflected by the tradeoff between FLZ and H_2_O_2_ resistance seen over time. These data explain the persistence of a heterogeneous fungal population and underscores the complexity and parallelism of evolution that is possible in the human lung during chronic disease.

## Results

### Naturally evolved *C. lusitaniae MRR1* alleles confer altered Mrr1 activity and FLZ resistance

Each of the twenty closely-related *C. lusitaniae* isolates from a single individual contained at least one nonsynonymous single nucleotide polymorphism (SNP) or single nucleotide insertion or deletion (indel) in *MRR1* relative to the deduced *MRR1* sequence of their most recent common ancestor (*MRR1*^*ancestral*^) (Fig. 1A) (10). To determine the impact of specific mutations in *MRR1* on Mrr1 activity, we expressed different *MRR1* alleles in a common genetic background in which the native *MRR1* had been deleted (U04 *mrr1*Δ). Deletion of *MRR1* in the FLZ-resistant strain U04 reduced the FLZ minimum inhibitory concentration (MIC) from 32 μg/ml to 4 μg/ml (10) and the decrease in MIC was complemented by restoring the native *MRR1*^*Y813C*^ allele (Fig. 1B). Complementation of U04 *mrr1*Δ with the *MRR1*^*ancestral*^ allele led to a FLZ MIC of 1 μg/ml which was 4-fold lower (*P*<0.0001) than the FLZ MIC of U04 *mrr1*Δ, suggesting that *MRR1*^*ancestral*^ had a function that reduced the FLZ MIC (Fig. 1B). Expression of an *MRR1* allele from a FLZ-sensitive isolate in the population (*MRR1*^*L1191H+Q1197\**^) also reduced the FLZ MIC to levels comparable to those for *MRR1*^*ancestral*^ (0.5-1 μg/ml) (Fig. 1B). Similar correlations between *MRR1* allele and FLZ MIC were observed when the *MRR1*^*ancestral*^, *MRR1*^*Y813C*^, and *MRR1*^*L1191H+Q1197\**^ alleles were expressed in a *mrr1*Δ derivative of the FLZ-sensitive strain U05, which indicated that strain background did not contribute to the FLZ MIC conferred by different *MRR1* alleles (Fig. 1C). We previously published that FLZ resistance correlated with expression of *MDR1* (10), also referred to as *MFS7* (19). Deletion of *MDR1* reduced the MIC, and the MIC was even lower in U04 *mrr1*Δ *mdr1*Δ (Fig. S1A) indicating that the moderately higher levels of FLZ resistance in U04 *mrr1*Δ compared to a strain with *MRR1*^*ancestral*^ was *MDR1*-dependent.

**Fig. 1:**
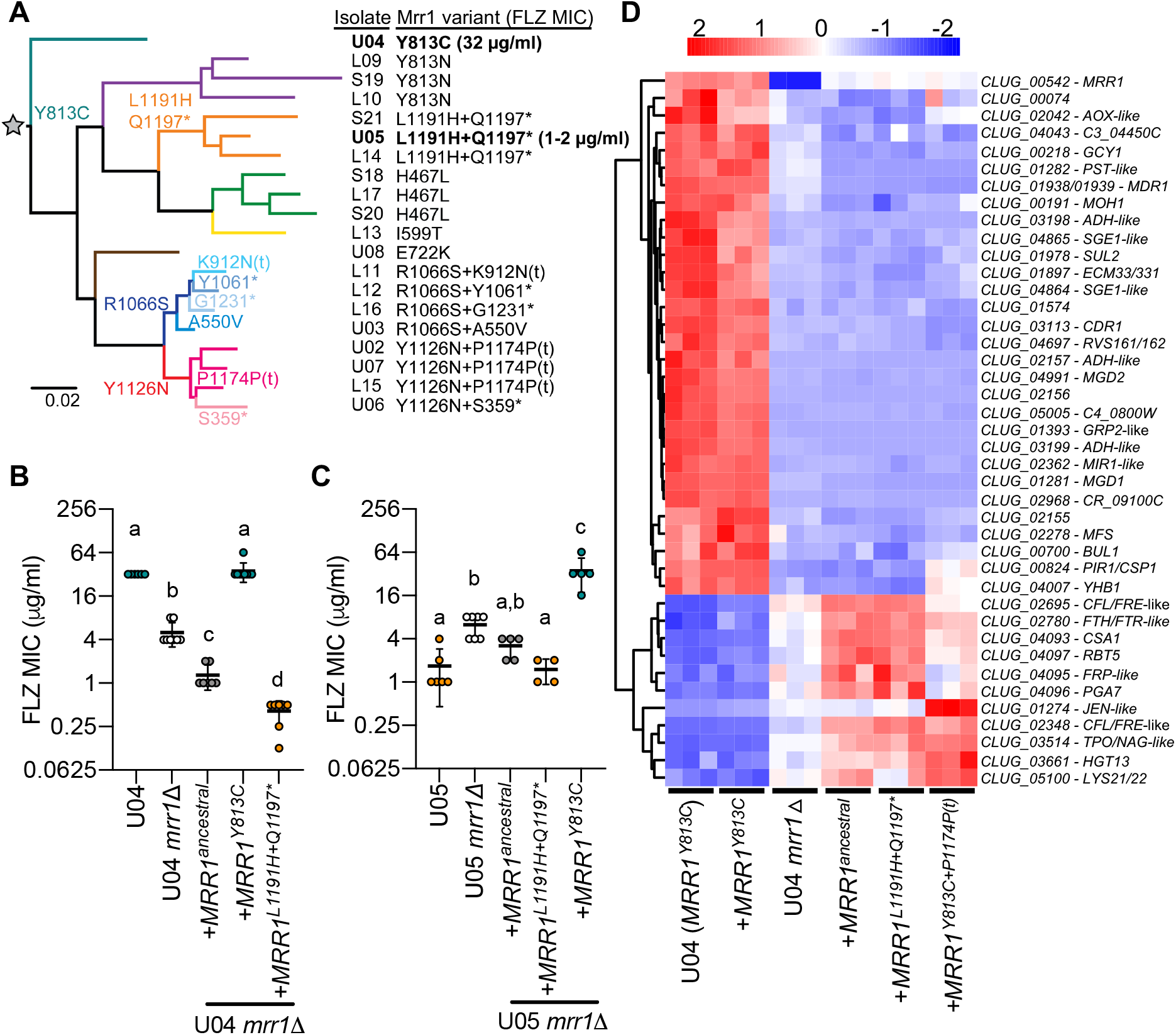
Naturally-evolved *MRR1* alleles confer higher or lower FLZ resistance compared to strains expressing the ancestral *MRR1* allele. **A**, Maximum likelihood-based phylogeny based on whole genome sequences of twenty previously sequenced *C. lusitaniae* isolates, modified from Demers, *et al.* (10). Select branchpoints are marked with the Mrr1 variants present in subsequent isolates. Mrr1 variants are identified by amino acid changes that resulted from SNPs or indels; * indicates a stop codon. The one nucleotide indel in codons P1174 (insertion) and K912 (deletion) cause frameshift mutations that resulted in early termination, denoted with ‘t’, at N1176 and L927, respectively. Gray star at the root of the tree denotes the ‘ancestral’ *MRR1* sequence, which lacks any of the mutations listed. **B**, FLZ MIC for unaltered, *mrr1*Δ or *MRR1* complemented strains in the FLZ-resistant U04 (native allele *MRR1*^*Y813C*^) strain background. **C**, Same as in **B**, but in the FLZ-sensitive U05 strain background (native allele *MRR1*^*L1191H+Q1197\**^). Strains containing the same *MRR1* allele in **B** and **C** are represented by circles of the same color. Data shown represents at least four independent assays on different days. One-way ANOVA with Tukey’s multiple comparisons test of log_2_ transformed values for **B**: all pairwise comparisons, *P*<0.0001 and **C**: all pairwise comparisons, *P*<0.001. **D**, Heatmap of normalized counts per million (CPM) from RNA-Seq analysis for genes that were differentially regulated between both strains expressing *MRR1*^*Y813C*^ (U04 and U04 *mrr1Δ* + *MRR1*^*Y813C*^) and U04 *mrr1*Δ + *MRR1*^*ancestral*^, U04 *mrr1*Δ *+ MRR1*^*L1191H+Q1197\**^, and U04 *mrr1*Δ when grown in liquid YPD medium. Complemented strains are denoted as by their respective *MRR1* alleles. Hierarchical clustering of row (genes) by Euclidean distance, cutoffs used were FDR <0.05 and fold change ≥2; additional information available in Table S1.

RNA-sequencing (RNA-seq) analysis validated the previously published result that *MDR1* correlated with the FLZ MIC for the different Mrr1 variants (10). The expression of *MGD1* and *MGD2*, two *C. lusitaniae* genes shown to be Mrr1-regulated, correlated with the expression of *MDR1* (Fig. 1D and Table S1) (10, 13, 33). Gene expression differences between U04 (*MRR1*^*Y813C*^), U04 *mrr1*Δ, and U04 *mrr1*Δ +*MRR1*^*Y813C*^ found that *mrr1*Δ is fully complemented upon return of *MRR1*^*Y813C*^ to the native locus (Fig. 1D and Fig. S2A for correlation plot) and that Mrr1 positively and negative regulates a large set of genes. Furthermore, a correlation analysis found that U04 *mrr1*Δ +*MRR1*^*ancestral*^ and U04 *mrr1*Δ +*MRR1*^*L1191H+Q1197\**^ were similar to each other but distinct from the *mrr1*Δ (Fig. S2A and 1B). A linear model comparing these strains identified forty-one genes with at least a 2-fold change in expression and corrected *P* value <0.05 (FDR). Comparison of non-isogenic *C. lusitaniae* strains similarly identified at least fourteen of the genes in Table S1 as putatively Mrr1-regulated (10, 13). Eighteen genes were homologs or had similar predicted functions as genes previously published as regulated by *C. albicans* Mrr1 (29), including *MDR1, FLU1* and multiple putative methylglyoxal reductases encoded by *GRP2*-like genes, such as *MGD1* and *MGD2* (Fig. 1D and Table S1). Other genes within the Mrr1 regulon are discussed further below.

The unexpected finding that FLZ MIC was higher upon deletion of *MRR1* relative to a strain with *MRR1*^*ancestral*^ or an allele from a FLZ-sensitive strain was also observed in distantly-related *C. lusitaniae* strains, ATCC 42720 and DH2383 (FLZ MICs of ~1-2 μg/ml). In both cases, deletion of *MRR1* led to a 2-4-fold increase in FLZ MIC to 4-8 μg/ml (Fig. S1B, *P*<0.001). The increase in FLZ MIC in *mrr1*Δ strains was not due to introduction of the selectable marker, *NAT1*, which encodes a nourseothricin acetyltransferase (34), as expression of *NAT1* from an intergenic site in the FLZ-sensitive U05 strain did not increase the FLZ MIC (Fig. S1C). These data led us to hypothesize that some Mrr1 variants (*MRR1*^*Y813C*^) lead to high Mdr1 activity while other Mrr1 variants (both *MRR1*^*ancestral*^ and the recently-diverged *MRR1*^*L1191H+Q1197\**^ alleles) repressed the expression of at least some Mrr1-controlled genes, such as *MDR1*. Indeed, the RNA-Seq analysis identified six genes, including *MDR1*, that while positively regulated when Mrr1 was constitutively active, were more highly expressed in U04 *mrr1*Δ than those strains encoding low activity Mrr1 variants (Fig. 1D and S2B). These data suggest that, for a small subset of Mrr1-regulated genes, including *MDR1*, low activity Mrr1 variants directly or indirectly inhibit gene expression.

### Truncation of *MRR1* has varied effects on Mrr1 activity and inducibility in clinical isolates

All twenty sequenced clinical *C. lusitaniae* isolates from a single human subject (Fig. 1A) had *MRR1* alleles with either one or two nonsynonymous mutations relative to *MRR1*^*ancestral*^, and we found that *C. lusitaniae* isolates with two mutations in *MRR1* had a significantly lower average FLZ MIC than isolates with a single *MRR1* mutation (Fig. 2A, *P*<0.001) (10). Interestingly, six of the seven *MRR1* alleles in the “two mutation” set encoded premature stop codons, resulting in loss of 34-906 amino acids (Fig. 2B). There were two instances in which the same mutation was found with different nonsense mutations (*) or single nucleotide indels that led to early termination (t): *MRR1*^***Y1126N***+*P1174P(t)*^ or *MRR1*^***Y1126N***+*S359\**^, and *MRR1*^***R1066****S*+*K912N(t)*^, *MRR1*^***R1066S***+*Y1061\**^ or *MRR1*^***R1066S***+*G1231\**^ (common mutation in bold, Fig. 1A) suggesting a complex evolutionary history for these alleles.

**Fig. 2:**
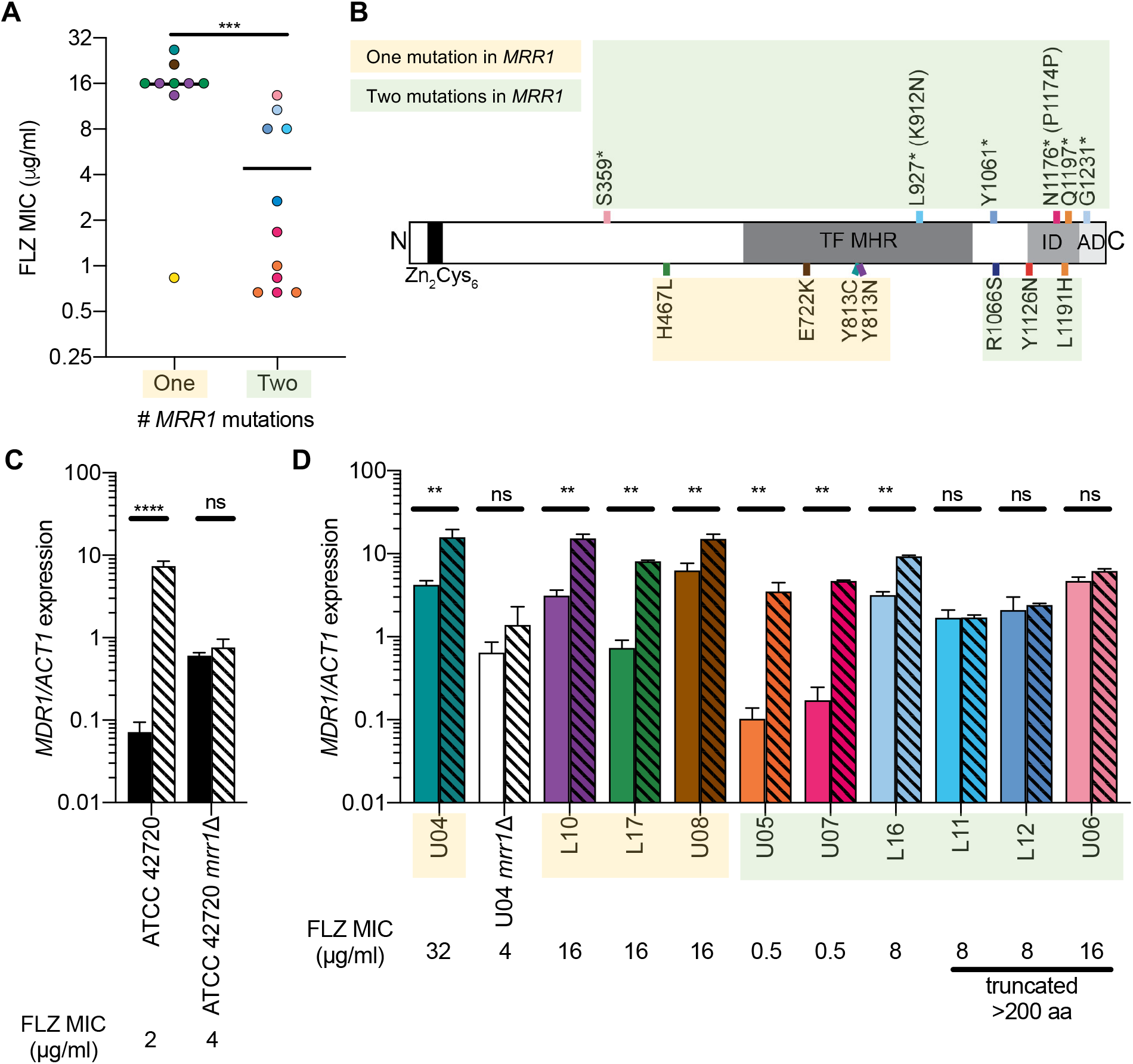
Premature stop codons in Mrr1 differentially impact *MDR1* induction by benomyl. **A**, Mean FLZ MIC for each of the twenty clinical *C. lusitaniae* isolates in Fig. 1A separated by the number of nonsynonymous mutations within *MRR1* (10); datapoints colored to match Fig. 1A, 2B and 2D. Mean of each group shown. Two-tailed unpaired t-test of log_2_ transformed MIC values; ***, *P*<0.001. **B**, Schematic of *C. lusitaniae MRR1* annotated with putative regulatory domains determined by sequence analysis or homology to *C. albicans* (22) and locations of truncating (above) and activating (below) mutations, colored to match Fig. 1A, 2A and 2D. Putative domains include the DNA binding domain with a zinc cluster motif (Zn_2_Cys_6_; amino acids 33 to 61), the transcriptional regulatory middle homology region (MHR; amino acids ~607-1023), an inhibitory domain (ID; amino acids 1123 to 1217) and an activating domain (AD; amino acids 1218 to 1265). L927 and N1176 are the locations of stop codons caused by indels in codons K912 and P1174, respectively. **C** and **D**, *MDR1* expression normalized to *ACT1* in the absence (solid) or presence (striped) of 50 μg/ml benomyl. Mean ± SD of representative data in biological triplicate shown, similar trends observed on at least three different days. In **D**, Bars are colored to correspond to Fig. 1A, and strains names are highlighted to correspond to the number of nonsynonymous SNPs in *MRR1*, yellow for one and green for two. **C** and **D**, Two-way ANOVA with Sidak’s multiple comparisons test; **, *P*<0.0; ****, *P*<0.0001; ns, not significant.

To better understand the effects of *MRR1* mutations on Mrr1 activity, we analyzed the effects of a chemical inducer of Mrr1 activity, benomyl (35–37), on *MDR1* expression. Benomyl strongly induced *MDR1* expression in an Mrr1-dependent manner in the FLZ-sensitive strain ATCC 42720 (Fig. 2C) and, to a lesser extent, in the FLZ-resistant strain U04, which has high basal *MDR1* expression (Fig. 2D) (10). Quantitative RT-PCR analysis of *MDR1* expression and induction by benomyl in this collection of clinical isolates with different Mrr1 variants found that the two isolates with the lowest basal *MDR1* expression and lowest FLZ MIC (U05 and U07) had the greatest induction by benomyl (34- and 27-fold, respectively) (Fig. 2D). Three isolates, L11, L12 and U06, had intermediate FLZ MICs and *MDR1* expression levels, and did not show benomyl induction, similar to *mrr1*Δ, and all encoded Mrr1 variants lacking greater than 200 amino acids leading us to propose that these mutations caused a loss of Mrr1 function (Fig. 2D). Other isolates showed a correlation between higher basal *MDR1* levels and elevated FLZ MICs, and this pattern was associated with lower relative levels of benomyl induction (Fig. 2D).

### Premature stop codons repeatedly arose in constitutively active Mrr1 variants and caused either a loss of constitutive Mrr1 activity or complete loss of function

In light of the mixed effects that these two-mutation *MRR1* alleles had on Mrr1 activity, we sought to determine the individual effects of mutations within each allele with a focus on the two strains with the lowest basal *MDR1* expression and the strongest induction of *MDR1* in response to benomyl: *MRR1*^*L1191H+Q1197\**^ (in U05) and *MRR1*^***Y1126N***+*P1174P(t)*^ (in U07) (Fig. 3A and 3B). We found that the *MRR1*^*L1191H*^ mutation caused a 32-fold increase in FLZ MIC (Fig. 3C) and 22-fold increase in *MDR1* expression (Fig. 3D) compared to *MRR1*^*ancestral*^ indicating that, like the Mrr1-Y813C variant, Mrr1-L1191H was constitutively active. In contrast, *MRR1*^Q1197*^, which caused the loss of 68 amino acids from the C-terminus of Mrr1, did not significantly alter the FLZ MIC compared to *MRR1*^*ancestral*^ allele indicating that it was neither a constitutively activating nor a null mutation (Fig. 3C). The reintroduction of the Q1197*\** mutation into *MRR1*^*L1191H*^, yielding *MRR1*^*L1191H+Q1197\**^, resulted in a 128-fold decrease in FLZ MIC (Fig. 3C) and 38-fold lower *MDR1* expression values (Fig. 3D) compared to a strain expressing *MRR1*^*L1191H*^ and led to a phenotype that mirrored that of *MRR1*^*ancestral*^. Benomyl inducibility of these variants is discussed below.

**Fig. 3:**
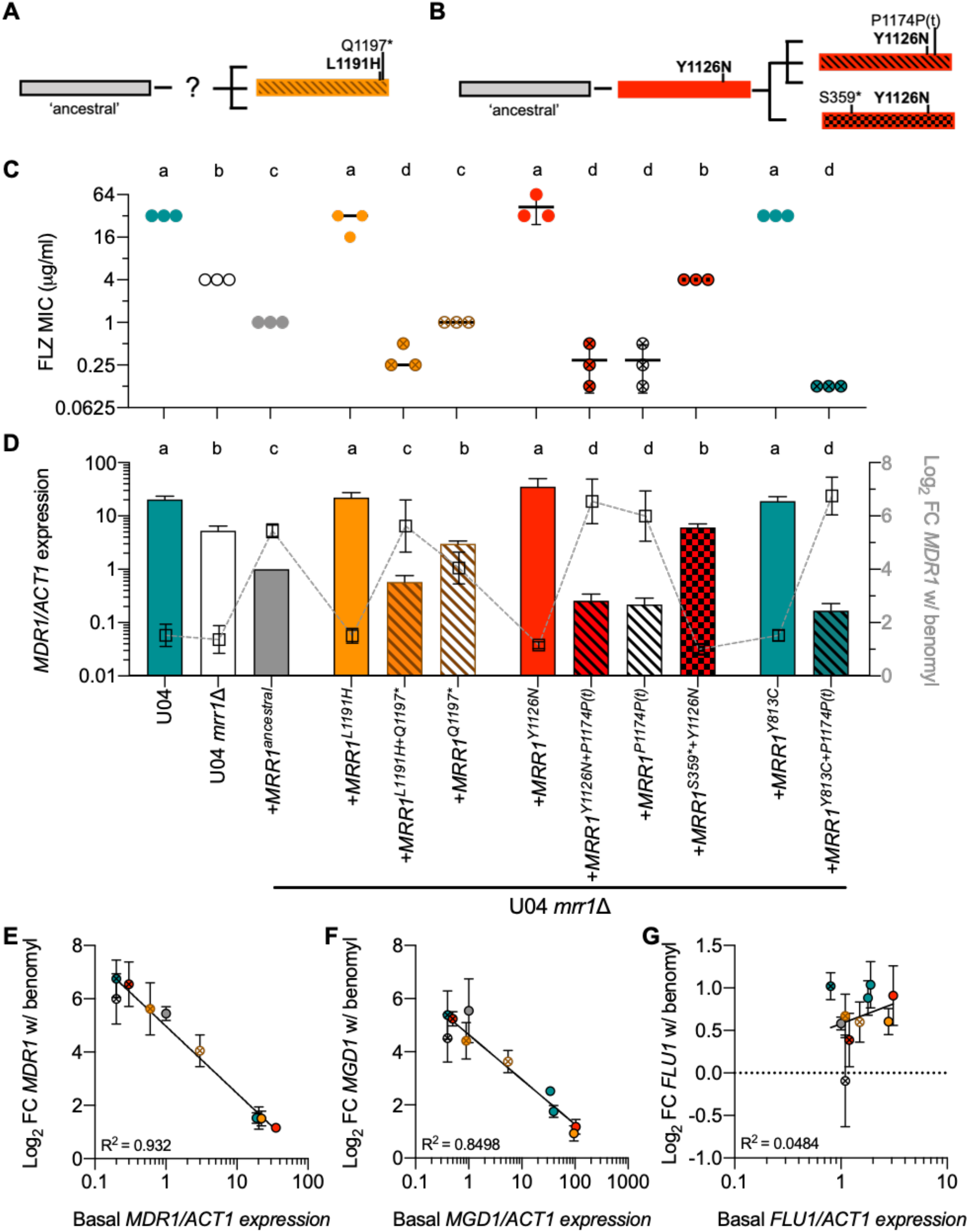
Premature stop codons repeatedly arose in constitutively active Mrr1 variants resulting in reduced basal Mrr1 activity, but in some cases restored Mrr1 inducibility. **A** and **B**, Schematic of inferred evolution of *MRR1* alleles in the L1191H+Q1197* (**A**) and Y1126N (**B**) lineages. **C**, FLZ MIC for U04, U04 *mrr1* Δ and *MRR1* complemented strains in the U04 *mrr1* Δ background, denoted by allele. Mean ± SD of three independent assays on different days shown. One-way ANOVA with Tukey’s multiple comparisons test of log_2_ transformed values; all pairwise comparisons, *P* <0.01. **D**, *MDR1* expression normalized to *ACT1* from culture grown in YPD (bars, left y-axis). Mean ± SD of three independent assays on different days; data from each day was normalized to the expression of U04 *mrr1* Δ + *MRR1*^*ancestral*^. One-way ANOVA with Tukey’s multiple comparisons testing of log_2_ transformed data; b-d, *P* <0.05; all other pairwise comparisons, *P* <0.01. Overlaid with log_2_ mean ± SD fold change in normalized *MDR1* expression following exposure to 50 μg/ml benomyl (squares, right y-axis); full data presented with statistics in Fig. S3D. **C** and **D**, FLZ MIC and *MDR1/ACT1* expression data are colored to match; the sample names are shown on the x-axis of Fig. 3D. **E-G**, Comparison of mean basal *MDR1* (**E**), *MGD1* (**F**) or *FLU1* (**G**) expression from Fig. 3D, S3B or S3C, excluding strains lacking functional *MRR1,* and mean ± SD log_2_ fold change (FC) induction following benomyl exposure from Fig. S3D-F. Goodness of fit r squared value for nonlinear regression shown.

*MRR1*^***Y1126N***+*P1174P(t)*^ (from U07) and *MRR1*^***Y1126N***+*S359\**^ (from the closely-related U06, Fig. 1A), were similarly analyzed (Fig. 3B). Expression of *MRR1*^*Y1126N*^ in U04 *mrr1*Δ created a strain with a high FLZ MIC (32-64 μg/ml, Fig. 3C) and *MDR1* expression (Fig. 3D), similar to that for strains with *MRR1*^*Y813C*^ or *MRR1*^*L1191H*^. Addition of the frameshift-inducing indel at P1174, which causes a premature stop codon at N1176 removing 89 amino acids, yielding *MRR1*^*Y1126N+P1174P(t)*^, caused a 128-fold decrease in the FLZ MIC and >100-fold decrease in *MDR1* expression relative to the strain expressing *MRR1*^*Y1126N*^ again leading to a strain that phenocopied that with *MRR1*^*ancestral*^ (Fig. 3C and 3D). The addition of the indel at P1174 into an allele with a different constitutively active variant, Mrr1-Y813C, (*MRR1*^*Y813C+P1174P(t)*^) also caused a 256- and >100-fold decrease in FLZ MIC and *MDR1* expression, respectively, (Fig. 3C and 3D). In contrast, addition of a SNP causing an early stop codon at S359 to the allele with the activating Y1126N mutation (*MRR1*^*Y1126N+S359\**^) yielded a strain that phenocopied U04 *mrr1*Δ, indicating this variant was inactive (Fig. 3C and 3D). Together, these data suggest that the Y1126N mutation caused constitutive Mrr1 activity, that was subsequently suppressed by premature stop codons that either restored Mrr1 repression of *MDR1* (P1174P(t)) or eliminated activity (S359*). The RNA-Seq analysis supported the results that premature stop codons near the very end of the protein converted constitutively active variants into ones that yielded expression profiles to those for *MRR1*^*ancestral*^ and that were distinct from *mrr1*Δ (Fig. 1D).

In addition to the differences in basal activity, the individual mutations alone and in combination affected chemical inducibility by benomyl. Levels of *MDR1* were strongly induced by benomyl in U04 *mrr1*Δ + *MRR1*^*ancestral*^ (40-fold increase), but not in the U04 parental strain with high Mrr1 activity or its *mrr1*Δ derivative (Fig. 3D). Along with the native Mrr1-Y813C, two other constitutively active Mrr1 variants (Mrr1-L1191H and Mrr1-Y1126N) showed only a 2-3-fold increase in *MDR1* expression with benomyl (Fig. 3D) similar to what was observed for more FLZ resistant clinical isolates (Fig. 2D). Surprisingly, addition of the mutations that caused premature stop codons within the last 100 amino acids of Mrr1 to the constitutively active Mrr1-L1191H, Mrr1-Y1126N and Mrr1-Y813C variants restored inducibility by benomyl (Fig. 3D). In fact, there was a strong and significant inverse correlation between basal *MDR1* expression and fold induction by benomyl (Fig. 3E).

As in *C. albicans*, *C. lusitaniae* Mrr1 regulates the expression of the methylglyoxal reductase encoded by *MGD1* (*CLUG_01281* or *GRP2*) (10, 29, 33, 38) and the multidrug efflux pump encoded by *FLU1* (*CLUG_05825*) (10, 39, 40) (Table S1 and Fig. S3A). As with *MDR1*, expression of both *MGD1* and *FLU1* was significantly higher in strains encoding the constitutively active Mrr1-Y813C, Mrr1-Y1126N and Mrr1-L1191H variants, compared to a strain encoding the Mrr1-ancestral variant, and the absence of the C-terminus in strains with activating mutations caused a significant decrease in basal *MGD1* and *FLU1* expression (Fig. S3B and S3C). Benomyl induction of *MGD1*, like *MDR1* (Fig. S3D), was restored upon loss of the C-terminus of the constitutively active Mrr1 variants further supporting the strong negative correlation between basal expression and induction by benomyl (Fig. 2F and S3E). *FLU1* expression, however, was not induced by benomyl in any strain suggesting that *FLU1* regulation by Mrr1 differs from *MGD1* and *MDR1* (Fig. 2G and S3F). Interestingly, *MDR1* and *MGD1*, while highly differentially expressed depending on Mrr1 activity (~20-fold or greater), were both de-repressed in the absence of Mrr1, and *FLU1* was not and was only weakly differentially expressed (<2-fold) (Fig. 1D and S2B). Together these data indicate the C-terminus of Mrr1 is required for constitutive expression of multiple Mrr1-regulated genes, but not for benomyl induction of the Mrr1-regulated genes tested (Fig. S3A). Combined with the Mrr1 activity across clinical isolates (Fig. 2D), these data indicate that in strains with constitutively active Mrr1 variants, there was selection for mutations to decrease Mrr1 activity, resulting in a mixed population containing constitutively active, truncated but inducible, and loss-of-function Mrr1 variants.

### Constitutive Mrr1 activity negatively impacts H_2_O_2_ resistance

We next sought to understand why mutations that reduce Mrr1 activity might repeatedly arise in this chronic infection. Previous studies have shown that overexpression of drug efflux pumps in drug resistant microbes can cause a fitness defect due to the energetic cost of constitutive pump production and activity in the absence of a selective substrate (41–43). Deletion of *MDR1* from U04 *mrr1*Δ +*MRR1*^*Y813C*^, which constitutively expresses *MDR1*, however, did not alter the growth rate across multiple carbon sources (Fig. 4A). In the absence of an obvious fitness defect, we considered factors present in the CF lung, which has been characterized as a highly inflamed environment containing an abundance of neutrophils and macrophages, and high oxidative stress (reviewed in (44, 45)). While little is known about the effects of fungus dominated chronic lung infections in CF, such as the infection from which these isolates originated, an analysis of cytokines within the bronchoalveolar lavage (BAL) fluid from the patient these isolates originated from showed pro-inflammatory cytokines (IL-8 and IL-1β) present were consistent with the neutrophilic environment seen in other patients with CF (Fig. 4B) (45).

**Figure 4:**
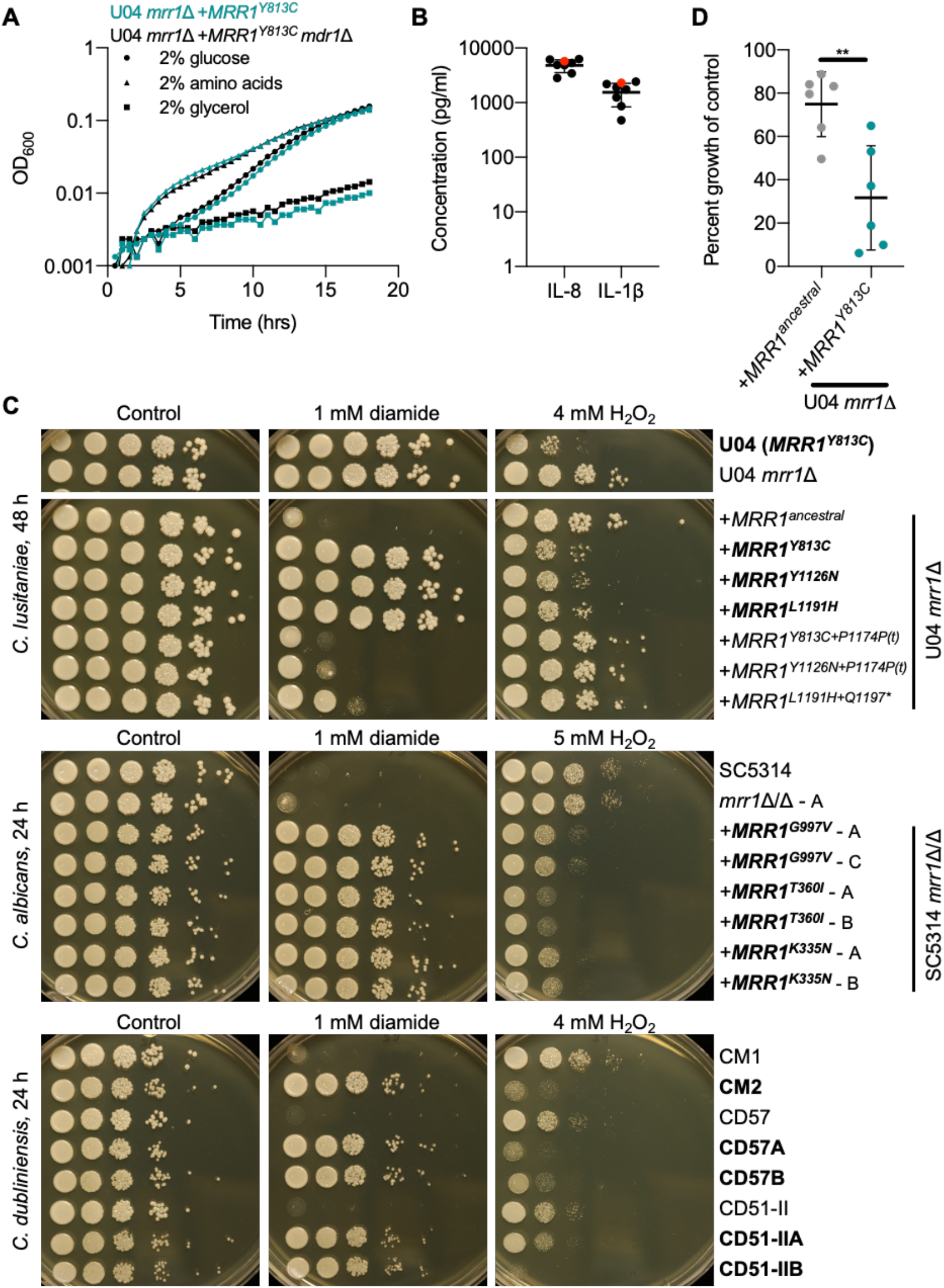
Tradeoff between constitutive Mrr1 activity conferring increased resistance to diamide but decreased resistance to hydrogen peroxide. **A**, Growth curve of U04 mrr1Δ + *MRR1*^*Y813C*^ (teal) and U04 *mrr1*Δ +*MRR1^Y813C^ mdr1*Δ (black) grown in YNB medium supplemented with the indicated carbon source: glucose (circles), amino acids (triangles), or glycerol (squares). Mean ± SD of representative data shown; grown at 37 °C. **B**, Quantification of cytokines IL-8 and IL-1β in BAL fluid from the CF patient with (red) or seven patients without (black) *C. lusitaniae* in their lungs. Two-way ANOVA with Sidak’s multiple comparisons testing found no significant differences. **C**, Serial dilution assays of *C. lusitaniae*, *C. albicans* and *C. dubliniensis strains* on YPD or YPD supplemented with the indicated concentration of diamide or H_2_O_2_. Strain names in bold have been shown in to contain GOF mutations in Mrr1 resulting in increased FLZ resistance (Fig. 3C and (40, 46, 47)). Plates were imaged after 24- or 48-hours growth at 37 °C, as indicated. **D**, Percent growth in well aerated 5 ml YPD + 1 mM H_2_O_2_ was calculated relative to the vehicle only control, after 22-24 hours growth at 37 °C. These data represent six independent assays performed on different days. Significance determined by paired t-test; \*\**P*<0.01.

In light of these findings, we investigated the effects of Mrr1 activity on reactive oxygen species (ROS) stress generated by hydrogen peroxide (H_2_O_2_), a stress strongly associated with high neutrophil counts. In a serial dilution assay, we found that isogenic strains encoding constitutively active Mrr1 variants, while highly resistant to FLZ and diamide (Fig. S4A), had increased sensitivity to 4 mM H_2_O_2_ compared to those expressing the Mrr1-ancestral variant (Fig. 4C). Diamide was used to illustrate relative Mrr1 activity instead of FLZ because serial dilution assays on rich medium (YPD) containing FLZ are not always representative of FLZ MIC, which are assessed in defined medium (Fig. S4B). Resistance to H_2_O_2_ was restored by addition of mutations causing both mild and severe premature stop codons (Fig. 4C, S4A). The effects of Mrr1 activity on H_2_O_2_ sensitivity were independent of strain background, as similar results were seen in isogenic strains in the U04 and U05 backgrounds (Fig. 4C and S4B). Surprisingly, deletion of *MDR1* from a strain encoding the constitutively active Mrr1-Y813C variant partially rescued growth (Fig. 4C and S4A), however, the absence of *MDR1* did not completely explain the differences as strains lacking *MRR1* had increased H_2_O_2_ resistance despite elevated *MDR1* expression (Fig. 4C). Additionally, the double mutant U04 *mrr1*Δ *mdr1*Δ did not have increased resistance to H_2_O_2_ compared to U04 *mrr1*Δ (Fig. 4C), suggesting this may be a complex response. A secondary assay quantifying growth after ~24 hours in liquid cultures containing 1 mM H_2_O_2_, though variable day-to-day, confirmed there was a reproducible difference in growth between strains encoding the low activity Mrr1-ancestral and constitutively active Mrr1-Y813C variants (Fig. 4D). Consistent with the plate-based assay, the absence of *MDR1* appeared to account for some but not all of the differences in growth in H_2_O_2_ (Fig. 4D). To determine if this phenomenon was unique to *C. lusitaniae* Mrr1 we examined a set of isogenic *C. albicans* isolates (40), and *in vivo* or *in vitro* evolved *C. dubliniensis* isolates (30) expressing *MRR1* alleles containing GOF mutations. We found that for all *C. albicans* and *C. dubliniensis* strain sets tested, strains with high Mrr1 activity, which were more resistant to FLZ (40, 46, 47) and diamide, were more sensitive to H_2_O_2_ than strains with low Mrr1 activity or lacking *MRR1* (Fig. 4C). These data show that the Mrr1 activity driven tradeoff between FLZ and H_2_O_2_ resistance is conserved across multiple *Candida* species.

A screen of isogenic strains for growth in varying concentrations of 48 chemical compounds resuspended from the Biolog Phenotype MicroArrays MicroPlates (Fig. S5) supported our findings that constitutive Mrr1 activity can increase sensitivity to oxidative stress. When comparing strains encoding either the low activity Mrr1-ancestral variant or the constitutively active Mrr1-Y813C variant, with either *MDR1* intact or removed, we found there were no differences in growth in the medium used to resuspend the Biolog compounds (Fig. S5A) and many conditions caused less than a 25% difference in growth (Fig. S5B). Unsurprisingly, constitutive Mrr1 activity conferred Mdr1-dependent resistance to twelve compounds, including three triazoles (FLZ, propiconazole, myclobutanil) (Fig. S5B and S5C). High Mrr1 activity also led to Mdr1-independent resistance to four additional compounds, including two other azoles (3-amino-1, 2, 4-triazole and miconazole nitrate) (Fig. S5B and S5C). Eight compounds caused a largely Mdr1-independent decrease in growth in strains encoding the constitutively active Mrr1-Y813C variant: 6-azauracil, berberine, BAPTA, lithium chloride, aminacrine, sodium metasilicate, pentamidine isethionate and potassium chromate (Fig. S5D). Interestingly, berberine and azaserine have previously been studied for their toxic effects on FLZ-resistant *Candida* strains (48, 49) and calcium inhibitors, such as BAPTA, have been reported to interfere with antifungal resistance (50, 51). While diverse, these compounds are broadly reported to effect metabolism and respiration (52–56), which can lead to oxidative damage via the production of ROS, and/or DNA/RNA integrity, either by direct binding or oxidative damage (57–62). Strain lacking *MRR1* or encoding a functional Mrr1 variant that contains a premature stop codon (<100 amino acids removed) were not sensitive to most of these compounds, suggesting secondary mutations causing a decrease or loss of Mrr1 activity could restore resistance in some environments (Fig. S5D).

To gain insight into the mechanisms that lead to differences in oxidative stress resistance between strains with different levels of Mrr1 activity, we compared the gene expression profiles after a 30-minute exposure to 0.5 mM H_2_O_2_, a partially inhibitory concentration. H_2_O_2_ exposure had broad strain-independent effects on the transcriptome, altering expression of 786 genes (FC≥2, FR<0.05) including increased expression of *CLUG_04072*, a homolog of *C. albicans CAT1*, which was previously shown to be important for the resistance of *C. lusitaniae* to H_2_O_2_ (63) (Fig. S6A and Table S2). While there were subtle differences in the H_2_O_2_ response between strains expressing the constitutively active Mrr1-Y813C variant compared to U04 *mrr1*Δ *MRR1*^*ancestral*^ there were no clear patterns that would explain the difference in H_2_O_2_ resistance (Fig. S6B). The majority of differences in gene expression were seen in the magnitude of induction of Mrr1-regulated genes by H_2_O_2_, a known inducer of Mrr1 in other species (29, 64), indicating that, as with benomyl (Fig. S3D-F), strains with constitutively active Mrr1variants are less inducible than strains with low activity variants (Fig. S6B and S6C). Next, we investigated the expression of homologs of oxidative stress response (OSR) genes previously characterized in *C. albicans* and *S. cerevisiae* and found that there was not a significant Mrr1-dependent difference in basal or H_2_O_2_-induced expression of these genes (Fig. S6A). Genes assessed included the oxidative stress responsive transcription factor encoded by *CaCAP1* or *ScYAP1*, superoxide dismutase (*SOD2, SOD4, SOD6*), enzymes involved in the thioredoxin (*TSA1, TRX1, TRR1*) and glutathione (*GPX, GSH1*) systems, catalase (*CAT1*), and OSR genes involved in carbohydrate metabolism and the DNA-damage response (65, 66). Further analysis is required to better understand the link between constitutive Mrr1 activity and H_2_O_2_ sensitivity, however these data highlight that the sensitivity is not due to failure to induce an oxidative stress response, but more-likely a consequence of the activity of Mrr1-regulated genes, such as *MDR1* (Fig. S4A and S4C).

### Phenotype dynamics in chronic infection populations over time

In light of the evidence for complex evolution of *MRR1* and the potentially advantageous phenotypes associated with both high and low Mrr1 activity, we sought to better understand the fractions of isolates with these Mrr1 associated traits over time. For this analysis, we used arrayed *C. lusitaniae* populations isolated from sputum or one BAL procedure collected from the same subject over three years, with the first time point approximately six months after the first clinical culture report of the high levels of “non-*albicans Candida*” (NAC) as shown in Fig. 5A. Upon plating isolates on agar with FLZ (8 μg/ml) or H_2_O_2_ (4 mM) (Fig. 5A), we found an inverse correlation between robust growth on FLZ and robust growth on H_2_O_2_. It was uncommon for isolates to be inhibited or uninhibited in both conditions (Fig. S5A). Isolates from the early samples were predominately sensitive to FLZ (10), but were largely resistant to H_2_O_2_. During and soon after the course of FLZ therapy (Sp1.5 and Sp2, respectively), however, there was an increase isolates that were more FLZ resistant but H_2_O_2_-sensitive (Fig. 5A). Subsequent samples from two years after the FLZ therapy was completed varied in the proportion of isolates that grew better on H_2_O_2_ and FLZ. Thus, the *C. lusitaniae* population shifted back and forth between being dominated by isolates with higher H_2_O_2_ resistance or higher FLZ resistance, but both phenotypes remained in the population over time (Fig. 5B).

**Fig. 5:**
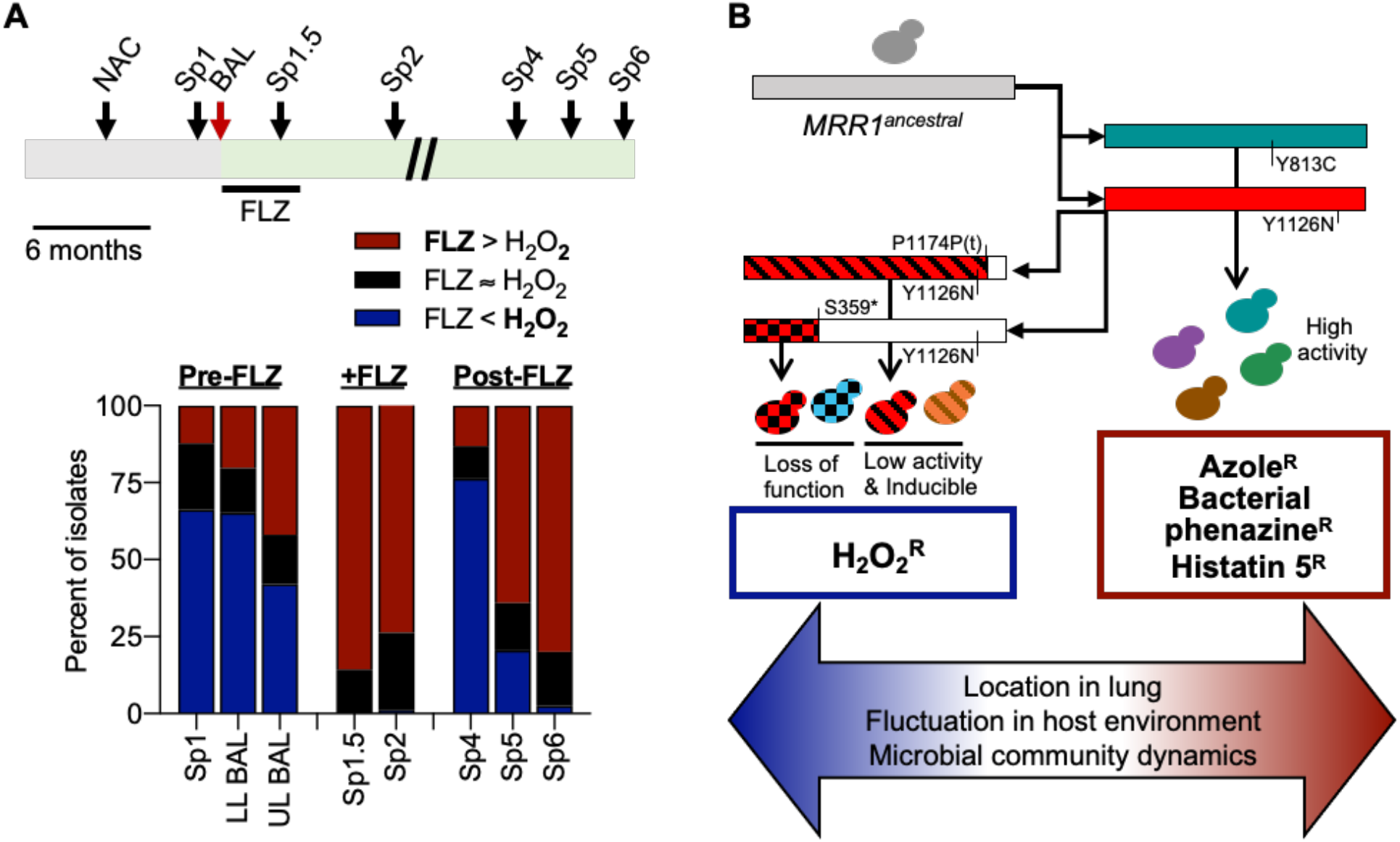
Tradeoff between FLZ and H_2_O_2_ resistance persists in evolving *C. lusitaniae* populations during a chronic lung infection. **A**, Schematic of sampling timeline (top) and histogram of the number of isolates that i) were mostly uninhibited on FLZ, but were inhibited by H_2_O_2_ (red), ii) were mostly uninhibited on H_2_O_2_ but were inhibited by FLZ (blue), or iii) grew similarly in both conditions (black). For the schematic, the gray bar represents the 6-10 months before the BAL during which this patient was identified as being colonized by non*-albicans Candida* (NAC) species. *C. lusitaniae* was determined to be the dominate microbe in the upper and lower lobe (UL and LL, respectively) BAL samples, which marks the start of the green bar. Sp1 was obtained one month before the BAL and was retrospectively also found to contain abundant *C. lusitaniae*. Sp1.5, Sp2, Sp4, Sp5 and Sp6 were obtained three, nine, thirty-two, thirty-five and thirty-eight months, respectively, after the BAL and all contained *C. lusitaniae.* A four-month course of FLZ therapy was given after the BAL. Scale of schematic is 1 inch = 6 months. Multiple isolates were collected from each sample/timepoint and assayed for growth on YPD supplemented with 8 μg/ml FLZ or 4 mM H_2_O_2_ after 48 hours at 37 °C. Growth was scored as completely inhibited, partially inhibited or uninhibited compared to a YPD only control. **B**, Model for the evolution of *C. lusitaniae MRR1* in this population. Whole genome sequencing and mutation analyses have shown that following the initial infection with *C. lusitaniae* encoding the Mrr1-ancestral variant, a combination of exposure to different stimuli that changed overtime or by locations within the CF lung environment lead to the selection for multiple constitutively active Mrr1 variants, some of which persisted over time and others that were subsequently mutated again resulting in premature stop codons that resulted in reversion to low activity that was inducible or complete loss of Mrr1 activity. The balance between selective pressures resulted in a heterogeneous population of isolates with varying resistance to biologically and clinically important compounds.

## Discussion

A population of *C. lusitaniae* isolates first described in Demers *et al.* (10) contained an unexpectedly large number of nonsynonymous mutations in the gene encoding the transcription factor, Mrr1, which regulated FLZ resistance, suggesting that Mrr1 activity was under strong selective pressure *in vivo*. These *MRR1* alleles contained either one or two nonsynonymous SNPs or indels (Fig. 1A) and isolates with one mutation had on average higher FLZ resistance than those with two nonsynonymous *MRR1* mutations (Fig. 2A). While multiple studies have shown that constitutive Mrr1 activity is beneficial under multiple biologically relevant conditions, including exposure to azoles (10, 29), bacterial-produced toxins including phenazines (10), and host-produced antifungal peptides including histatin 5 (10, 40), it was unclear why *MRR1* alleles conferring low Mrr1 activity would be selected for in this population (10). Deconstruction of *MRR1* alleles with two mutations revealed an evolutionary path on which an activating mutation arose first, followed by suppressing mutations that either restored low basal activity but retained inducibility, or abolished Mrr1 activity altogether (Fig. 3 and S3). Interestingly, a *C. parapsilosis* strain was recently found to contain a central domain mutation and a C-terminal truncation (Mrr1^P295L+Q1074*^) similar to the alleles described above, however, it is not currently known how these mutation impact Mrr1 activity and FLZ resistance (67), suggesting that selection for and against elevated Mrr1 activity may also occur in other *Candida* species.

Surprisingly, the RNA-Seq analysis of isogenic strains expressing different *MRR1* alleles revealed that *C. lusitaniae* Mrr1 appears to positively and negatively regulate genes expression (Fig. 1D) although further analysis is required to determine which genes are direct targets of Mrr1. Adding to previous studies in *C. lusitaniae* (10, 13) and *C. albicans* (29), we found that Mrr1 positively regulates 41 genes with a fold change ≥ 2 and 102 genes with a fold change ≥ 1.5 (FDR<0.05). Mrr1-induced genes include multiple MFS and ABC transporters (*i.e. MDR1, FLU1, CDR1*), methylglyoxal reductases (33), putative alcohol dehydrogenases, and a variety of other putative metabolic genes (Table S1). Constitutively active Mrr1 also appears to repress expression of 42 genes (fold change ≥ 1.5, FDR<0.05), including multiple iron and/or copper transporters and reductases, and sugar transporters (Table S1). These data combined with Bierman *et al.*, which showed that *C. lusitaniae* Mrr1 is induced by the spontaneously formed stress signal methylglyoxal (33), imply that Mrr1 may play a larger role in a generalized metabolic or stress response, beyond what has been previously studied in response to FLZ and xenobiotic stressors.

While the C-terminal region of *C. lusitaniae* Mrr1 was necessary for constitutive Mrr1 activity, it was not required for induction of Mrr1-regulated genes, including *MDR1* and *MGD1*, in response to benomyl (Fig. 2D, 3 and S3). Subsequent addition of mutations resulting in the loss >200 amino acids, however, caused a slight decrease in FLZ resistance and *MDR1* expression, but these variants were no longer inducible by benomyl and phenocopied strains completely lacking *MRR1* (Fig. S1B, 3C and 3D). These data are consistent with previous studies showing C-terminal truncations prior to amino acid 944 in *C. albicans MRR1*, homologous to position 1116 in *C. lusitaniae MRR1*, caused a complete loss of CaMrr1 activity (68). The L11, L12 and U06 strains encoding Mrr1 variants with premature stop codons before amino acid 1116 similarly phenocopied *mrr1*Δ strains (Fig. 2C and 2D). Surprisingly, loss-of-function Mrr1 variants and *mrr1*Δ strains had intermediate expression of a subset of the most strongly differentially regulated genes compared to strains with low activity Mrr1 (Fig. S2B), which has not been observed in other *Candida* species (29, 31, 32). Additional studies are required to determine if this phenomenon is unique to *C. lusitaniae* or more broadly shared among non-*albicans Candida* species closely related to *C. lusitaniae*, such as *C. auris* (20, 26), and if any of the co-regulators of the Mrr1 regulon described in *C. albicans* (64, 69, 70) are involved. These findings raised the question as to why, if constitutive Mrr1 was initially selected for, would it later be selected against *in vivo*, especially in the absence of an obvious growth defect (Fig. 4A and S5A).

Chronic lung infections are typically an inflamed environment (Fig. 4B) containing a high number of polymorphonuclear leukocytes (PMNs) that produce proteases, myeloperoxidases and ROS (71, 72), which is an important component of the immune system used to kill fungi (reviewed in (73)). In a screen of diverse chemical compounds, we found that strains with constitutive Mrr1 activity were more strongly inhibited by multiple compounds that have previously been shown to cause damage through oxidative stress (Fig. S5B-D). When we specifically interrogated H_2_O_2_ resistance, we found that *C. lusitaniae* strains encoding constitutively active Mrr1 variants were more sensitive than strain encoding low activity Mrr1 variants or lacking a functional Mrr1 (Fig. 4A, S4). Sensitivity to H_2_O_2_ and the compounds from the Biolog plates was at least partially dependent on Mdr1, thought other Mrr1-regulated genes may still contribute to the decreased growth under conditions of oxidative stress (Fig. S4B, S5 and S6). Interestingly, the tradeoff between FLZ and H_2_O_2_ resistance was conserved broadly among a time series of *C. lusitaniae* isolates and other *Candida* species (Fig. 4C and 5A).

As outlined in the model in Figure 5B, together these data highlight that changing environments within complex and dynamic chronic infections could contribute to the development of heterogeneous fungal populations. Though it appears that initial selection on the ancestral version of Mrr1 was driven by the need for increased Mrr1 activity, over time either these selective pressures were removed, or other pressures became dominant, resulting in a secondary wave of mutations. This secondary wave of mutations caused a decrease or loss of Mrr1 activity, which uniquely to *C. lusitaniae* further contributed towards a population with mixed levels of FLZ resistance (Fig. 5B). Though the exact selective pressures at play in this instance are unknown, these data highlight the importance of understanding how microbes evolve *in vivo*, as complex environments, even in the absence of clinically used antifungals, can shape the microbial population and lead to antimicrobial resistance.

## Materials and Methods

### Strains and growth conditions

*Candida* strains used in this study are listed in Table S3. All strains were stored as frozen stocks with 25% glycerol at −80 °C and subcultured on YPD (1% yeast extract, 2% peptone, 2% glucose, 1.5% agar) plates at 30 °C. Strains were regularly grown in YPD liquid medium at 30 °C on a roller drum. Cells were grown in YNB (0.67% yeast nitrogen base medium with ammonium sulfate (RPI Corp)) liquid supplemented with either 2% glucose, 2% glycerol or 2% casamino acids and in RPMI-1640 (Sigma, containing L-glutamine, 165 mM MOPS, 2% glucose) liquid as noted. Media was supplemented with 8 μg/ml FLZ (stock 4 mg/ml in DMSO), 1 mM diamide (stock 58 mM in water) or 1-6 mM H_2_O_2_ (30% w/v in water, ~9.8M) as noted. *Escherichia coli* strains were grown in LB with either 100 μg/ml carbenicillin or 15 μg/ml gentamycin as necessary to obtain plasmids. BAL fluid and sputum were obtained in accordance with institutional review board protocols as described in (74).

### DNA for gene knockout constructs

Gene replacement constructs for knocking out *MRR1* (*CLUG_00542*, as annotated in (10)) and *MDR1* (*CLUG_01938/9* (10)) were generated by fusion PCR as described in Grahl *et al*. (63). All primers (IDT) used are listed in Table S4. Briefly, 0.5 to 1.0 kb of the 5’ and 3’ regions flanking the gene was amplified from U04 DNA, isolated using the MasterPure Yeast DNA Purification Kit (epiCentre). The nourseothricin (*NAT1*) or hygromycin (*HygB*) resistance cassette was amplified from plasmids pNAT (75) and pYM70 (76), respectively, using the Zyppy Plasmid Miniprep kit (Zymo Research). Nested primers within the amplified flanking regions were used to stitch the flanks and resistance cassette together. PCR products for transformation were purified and concentrated with the Zymo DNA Clean & Concentrator kit (Zymo Research) with a final elution in molecular biology grade water (Corning).

### DNA for insertion of *NAT1* at neutral site in *C. lusitaniae* genome

The approximately 4000 bp genomic region between *CLUG_03302* and *CLUG_*03303 on chromosome 4, which was not predicted to contain any genes or promoter regions, was targeted as a potentially neutral insertions site. To create plasmid DH3261 containing *NAT1* flanked by homology to this region of chromosome 4, approximately 1.0 kb of the flanking regions (positions 228,652 – 229,651 and 229,701 – 230,691) were amplified from U05 gDNA. All primers (IDT) used are listed in Table S4. *NAT1*was amplified from pNAT (75). PCR products were purified and concentrated then assembled with the vector (pRS426 (77) linearized with KpnI-HF and SalI-HF (New England BioLabs) and treated with the phosphatase rSAP (New England BioLabs)) using the NEBuilder HiFi DNA Assembly cloning kit (New England BioLabs). Assemblies were transformed into High Efficiency NEB®5-alpha competent *E. coli* (New England BioLabs). The *NAT1* insertion construct was isolated from DH3261 by digestion with KpnI-HF and SalI-HF (New England BioLabs).

### Plasmids for complementation of *MRR1*

Plasmids for complementing *MRR1* were created as described in Biermann *et al*, 2020 (33). For naturally occurring *MRR1* alleles, we amplified i) the *MRR1* gene and terminator with ~1150 bp upstream for homology from the appropriate strain’s genomic DNA, ii) the selective marker, *HygB* from pYM70 (76), and iii) ~950 bp downstream of *MRR1* for homology from genomic U05 (identical sequence for all relevant strains) using primers (IDT) listed in Table S4. PCR products were cleaned up using the Zymo DNA Clean & Concentrator kit (Zymo Research) and assembled using the *S. cerevisiae* recombination technique previously described (78). Plasmids created in *S. cerevisiae* were isolated using a yeast plasmid miniprep kit (Zymo Research) and transformed into High Efficiency NEB®5-alpha competent *E. coli* (New England BioLabs). *E. coli* containing pMQ30 derived plasmids were selected for on LB containing 15 μg/ml gentamycin. Plasmids from *E. coli* were isolated using a Zyppy Plasmid Miniprep kit (Zymo Research) and subsequently verified by Sanger sequencing with the Dartmouth College Genomics and Molecular Biology Shared Resources Core. *MRR1* complementation plasmids were linearized with Not1-HF (New England BioLabs), cleaned up the Zymo DNA Clean & Concentrator kit (Zymo Research) and eluted in molecular biology grade water (Corning) before transformation of 2 μg into *C. lusitaniae* strain U04 *mrr1*Δ as described below.

The *MRR1*^*ancestral*^ allele sequence was amplified from gDNA of a closely related *C. lusitaniae* isolate that had the same *MRR1* sequence but lacked any of the nonsynonymous mutation that varied among the population of *C. lusitaniae* isolates described here. This *MRR1* sequences does contain multiple synonymous and nonsynonymous mutations in comparison with that of the reference strains, ATCC 42720 (79). Additional *MRR1* alleles were amplified from gDNA from U04 (*MRR1*^*Y813C*^), U05 (*MRR1*^*L1191H+Q1197\**^), U02 (*MRR*^*Y1126N+P1174P(t)*^) and U06 (*MRR1*^*S359*+Y1126N*^). While making the pMQ30^*MRR1-S359*+Y1126N*^ plasmid, one clone was identified that lacked the S359* mutation resulting in the pMQ30^*MRR1-Y1126N*^ plasmid. To create additional *MRR1* alleles that were not identified within any *C. lusitaniae* isolates, pieces of *MRR1* were selectively removed and repaired with DNA either containing or lacking the desired mutation. Because the L1191H and Q1197* mutations were so close together, an alternate strategy was used to separate these mutations from the *MRR1*^*L1191H+Q1197\**^ allele. DNA fragments synthesized by IDT containing either the L1191H or Q1197* mutations alone (sequences in Table S4) were amplified then assembled with pMQ30^*MRR1-L1191H+Q1197\**^ (linearized with PvuI-HF) using the NEBuilder HiFi DNA Assembly cloning kit (New England BioLabs). To remove an unexpected nonsynonymous mutation in pMQ30^*MRR1-Q1197\**^, this plasmid was digesting with EcoNI and repaired with a piece of DNA amplified from U04 *mrr1*Δ+*MRR1*^*ancestral*^ lacking the unwanted mutation. pMQ30^*MRR1*^ complementation plasmids was digested with Not1-HF (New England BioLabs) for transformation.

### Strain construction

Mutants were constructed as previously described in Grahl *et al.* using an expression free ribonucleoprotein CRISPR-Cas9 method (63). 1 to 2 μg of DNA for gene knockout constructs generated by PCR or 2 μg of digested plasmid, purified and concentrated with a final elution in molecular biology grade water (Corning), was used per transformation. Plasmids containing complementation and knockout constructs and resulting strains are listed in Table S3 and crRNAs (IDT) are listed in Table S4. Transformants were selected on YPD agar containing 200 μg/mL nourseothricin or 600 μg/mL hygromycin B.

### Drug susceptibility assays

Minimum inhibitory concentration (MIC) was determined using a broth microdilution method as previously described (80) with slight modifications (10). Briefly, 2×10^3^ cells were added to a two-fold dilution series of FLZ prepared in RPMI-1640, starting at an initial concentration of 64 μg/ml, then incubated at 35 °C for 24 hours. The MIC was defined as the drug concentration that abolished visible growth compared to a drug-free control.

### Quantitative RT-PCR

Overnight cultures were back diluted to an OD_600_ of ~0.1 and grown for 6 hours in YPD liquid medium at 30°C. 50 μg/ml of benomyl (stock 10 mg/ml in DMSO) or an equivalent volume of DMSO were added for experiments assessing the induction of Mrr1 activity. 7.5 μg RNA (harvested using the MasterPure Yeast RNA Purification Kit (Epicentre)) was DNAse treated with the Turbo DNA-free Kit (Invitrogen). cDNA was synthesized from 300-500 ng of DNAse-treated RNA using the RevertAid H Minus First Strand cDNA Synthesis Kit (Thermo Scientific), following the manufacturer’s instructions for random hexamer primer (IDT) and GC rich template. qRT-PCR was performed on a CFX96 Real-Time System (Bio-Rad), using SsoFast Evergreen Supermix (Bio-Rad) with the primers listed in Table S4. Thermocycler conditions were as follows: 95 °C for 30 s, 40 cycles of 95 °C for 5 s, 65 °C for 3 s and 95 °C for 5 s. Transcripts were normalized to *ACT1* expression.

### RNA Sequencing

Overnight cultures were back diluted into YPD and grown to exponential (~8 h) twice, then treated with vehicle or 0.5 mM H_2_O_2_ for 30 minutes, in biological triplicate. RNA was harvested from snap-frozen pellets (using liquid nitrogen) using the MasterPure Yeast RNA Purification Kit (Epicentre) and stored at −80 °C. RNA libraries were prepared using the Kapa mRNA HyperPrep kit (Roche) and sequenced using single-end 75 bp reads on the Illumina NextSeq500 platform. The data analysis pipeline is available in github repository (https://github.com/stajichlab/RNASeq_Clusitaniae_MRR1) and archived as DOI: [To be generated]. FASTQ files were aligned to the ATCC 42720 (79) genome with the splice-site aware and SNP tolerant short read aligner GSNAP (v v2019-09-12) (81). The alignments were converted to sorted BAM files with Picard (v2.18.3; https://broadinstitute.github.io/picard/) and read counts computed with featureCounts (v1.6.2) (82) with updated genome annotation to correct truncated gene model for locus *CLUG_00542*, and combine a single gene split into two, *CLUG_01938_1939*; reasoning for these changes explained in (10). Differential gene expression analyses were performed with the edgeR (83) package in Bioconductor, by fitting a negative binomial linear model. The resulting *P* values were corrected for multiple testing with Benjamini-Hochberg to control the false discovery rate. Genes for which there were less than 2 counts per million (CPM) across the three (absent genes) were not included for differentially expressed gene analysis. Two separate linear models were created to define the Mrr1 regulon in control conditions alone and determine the interaction between Mrr1 activity and H2O2 exposure.

To define the Mrr1 regulon in YPD alone we identified genes differentially expressed between strains with constitutive Mrr1 activity (U04 and U04 *mrr1*Δ +*MRR1*^*Y813C*^) and low Mrr1 activity (U04 *mrr1*Δ, U04 *mrr1*Δ +*MRR1*^*ancestral*^, and U04 *mrr1*Δ +*MRR1*^*L1191H+Q1197\**^); this model contained 5,474 genes. We discarded genes for which i) the log2FC greater than 1 (2-fold, see Fig. 1B) or 0.585 (1.5-fold, see Table S1) with an FDR<0.05, (ii) the average CPMs for replicates was not greater than 10 for any strain, and ii) expression in both U04 and U04 *mrr1*Δ +*MRR1*^*Y813C*^ was similar. Results are summarized in Table S1, including the Mrr1 regulon (Table S1a), and the normalized CPMs/gene used for this linear model(Table S1b).

To determine how constitutive Mrr1 activity impacted the response to H_2_O_2_ we identified the overlap between the interaction between U04 or U04 *mrr1*Δ +*MRR1*^*Y813C*^ and exposure to 0.5 mM H2O2, as compared to the reference strain (U04 *mrr1*Δ +*MRR1*^*ancestral*^) and condition (YPD alone); this model contained 5600 genes. Results are summarized in Table S2, including the interaction between strains with constitutively active Mrr1 and H2O2 (Table S2a), the effect of H2O2 treatment (Table S2b), and all normalized CPMs/gene used for this linear model (Table S2c).

### Biolog Phenotype MicroArrays analysis

For the chemical sensitivity screen, the chemicals in Biolog plates PM22D and PM24C were resuspended in 100 ul YPD liquid and transferred to a sterile 96-well polystyrene plate (Fisher) for kinetic measurements. 100 ul of cells adjusted to an OD of 0.01 in YPD was added to each well. Plates were incubated at 37 °C for 24 hours. A control plate containing no drug was grown simultaneously for comparison.

### Serial dilution assays

Following growth in YPD medium overnight with aeration at 30°C, cultures were diluted in water to an OD_600_ of 1. Serial dilutions of ten-fold were carried out in a microtiter plate to yield six concentrations ranging from approximately 10^7^ cells/ml (for OD_600_ of 1) to approximately 10^2^ cells/ml. 5 μl of each dilution were applied to YPD plates containing 4 or 5 mM H_2_O_2_ (stock 30% w/v, 9.8 M) or 1 mM diamide (stock 58 mM in water). Images were captured after incubation at 37°C for 24 or 48 hours.

### Luminex Analysis

Cytokines in BAL fluid samples were measured (pg/ml) in singlicate by Luminex using a Millipore human cytokine multiplex kits (EMD Millipore Corporation, Billerica, MA) according to manufactures instructions. Assays were performed by the DartLab – Immune Monitoring and Flow Cytometry Resource core at Dartmouth.

### Statistical analyses

Statistical analyses were done using GraphPad Prism 6 (GraphPad Software). Unpaired Student’s t-tests (two-tailed) with Welch’s correction were used to evaluate the difference in FLZ MIC between isolates containing one of two mutations in *MRR1*. One and two-way ANOVA tests were performed across multiple samples with either Tukey’s multiple comparison test for unpaired analyses or Sidak’s multiple comparison test for paired analyses conducted in a pairwise fashion. *P* values <0.05 were considered as significant for all analyses performed and are indicated with asterisks: \**P*<0.05, \*\**P*<0.01, \*\*\**P*<0.001 and \*\*\*\**P*<0.0001.

## Data availability

The data supporting the findings in this study are available within the paper and its supplemental information and are also available from the corresponding author upon request. The raw sequence reads from the RNA-Seq analysis have been deposited into NCBI sequence read archive under BioProject PRJNA680763. Raw and processed RNA-Seq count data are available in Gene Expression Omnibus (GSE162151) and include minor updates to the genome annotation and assembly for *C. lusitaniae*.

## Acknowledgments

We would like to thank J. Morschhäuser (Universität Würzburg) and L. Myers (Dartmouth College) for sharing *C. albicans* and *C. dubliniensis* strains. Research reported in this publication was supported by National Institute of Health (NIH) grant R01 AI127548 to D.A.H. from the National Institute of Allergy and Infectious Disease, R01 HL122372 to A.A. from the National Heart, Lung and Blood Institute, and National Institute of General Medical Sciences (NIGMS) of the NIH under award number T32GM008704 and AI133956 to E.G.D. J.E.S. is a CIFAR Fellow in the program Fungal Kingdom: Threats and Opportunities. This work was also supported by the Cystic Fibrosis Foundation Research Development Program (CFFRDP) STANTO19R0 for the Translational Research Core, and ASHARE20P0 to A.A. Sequencing services and specialized equipment was provided by the Genomics and Molecular Biology Shared Resource Core at Dartmouth, and Luminex analysis was performed by the DartLab – Immune Monitoring and Flow Cytometry Resource Core at Dartmouth, both supported by NCI Cancer Center Support Grant 5P30CA023108-37. Equipment used was supported by the NIH IDeA award to Dartmouth BioMT P20-GM113132. Analyses were performed using the computational and data storage resources of the University of California-Riverside HPCC funded by grants from the National Science Foundation (NSF) (MRI-1429826) and NIH (1S10OD016290-01A1). The content is solely the responsibility of the authors and does not necessarily represent the official views of the NIH.

